# Continued topographical learning- and relearning-dependent activity in the resting state after post-training sleep and wake

**DOI:** 10.1101/2020.08.17.254383

**Authors:** Michele Deantoni, Thomas Villemonteix, Evelyne Balteau, Christina Schmidt, Philippe Peigneux

**Affiliations:** Neuropsychology and Functional Neuroimaging Research Unit (UR2NF) at CRCN - Center for Research in Cognition and Neurosciences and UNI - ULB Neurosciences Institute, Université Libre de Bruxelles (ULB), CP191 Av F Roosevelt 50, B1050 Bruxelles, Belgium; Psychopathology and Neuropsychology Lab, Paris 8 University, Rue de la Liberté 2, 93526 Saint-Denis, France; CRC-GIGA In Vivo Imaging Université de Liège, Allée du 6 Août, Bâtiment B30, Sart Tilman, 4000 Liège, Belgium; Psychology and Neurosciences of Cognition (PsyNCog), Quartier Agora, Place des Orateurs, 3, Bâtiment B33, 4000 Liège, Belgium

**Keywords:** functional MRI, sleep deprivation, memory consolidation, resting-state fMRI, ALFF, spatial learning.

## Abstract

Continuation of experience-dependent neural activity during offline periods of sleep and wakefulness is a critical element of memory consolidation. Using functional magnetic resonance imaging (fMRI), offline consolidation effects have been evidenced probing behavioral and neurophysiological changes during memory retrieval, i.e. in the context of task practice. Resting state fMRI (rfMRI) further allows investigating the offline evolution of recently learned information, without confounding task-related effects. In the present study, we used rfMRI to investigate offline changes in functional connectivity (FC) and Amplitude of Low Frequency Fluctuation (ALFF) associated with learning and relearning in a spatial navigation task, following an episode of post-training wake or sleep, respectively. Resting state activity was measured immediately (i) before and (ii) after topographical learning in a virtual town, (iii) 4 days later after regular sleep (RS) vs. sleep deprivation (SD) on the first post-learning night, and (iv) after topographical re-learning in an extended town encompassing the initial map. Task (navigation)-based fMRI activity was also recorded on Day 1 and Day 4 during target retrieval. Our results highlight the continuation of navigation-related activity in the subsequent resting state, as evidenced by changes in FC and ALFF in task-related neural networks. Behavioural performance was not modulated by post-training SD vs RS. However, in line with prior reports, post-training SD was associated with increased FC between navigation-related brain structures when faced to the task of learning a novel but related environment in the extended version of the city at Day 4. These results suggest the use of compensatory resources to link novel information with SD-related less efficiently consolidated memory traces.

## INTRODUCTION

Evidence ranging from cell recordings in rodents to behavioral and human neuroimaging data shows that sleep participates in the long-term consolidation of recently acquired information ^1^ in various memory domains ^2,3^. Amongst others, sleep supports the consolidation of spatial memory ^4–6^, the system responsible for recording information about one’s environment and spatial orientation. Spatial and episodic declarative memory share a similar neuroanatomical system in human beings and animals ^7^. This makes spatial memory an attractive paradigm to study the effects of sleep, with unique opportunities for translational inferences.

Neuroimaging studies consistently showed that spatial navigation in human beings is subtended by a wide brain network encompassing the hippocampus, the dorsal striatum, the precuneus, and the entorhinal, parahippocampal, retrospenial and frontal cortices ^8,9^. Rodent studies demonstrated that the hippocampus is involved in the rapid acquisition of spatial information at the early stage of learning, allowing the animal to reach its target from any starting position ^10^. The entorhinal cortex contributes to the encoding of map-like spatial codes, whereas parahippocampal and retrosplenial cortices provide significant inputs allowing to anchor these cognitive maps to fixed environmental landmarks. Spatial encoding- and retrieval-related activity in these brain areas is coordinated with frontal lobe activity to plan routes during navigation ^11^. Besides, navigation performance also rely on the dorsal striatum, which supports a complementary learning system based on rewarded stimulus-response automated behavioral associations, such as gradually learning successive body turns in response to environmental cues to reach a known target location from a starting point ^10,12^. Human and animal studies investigating the role of hippocampal and striatal areas in place-based vs. response-based navigation strategies suggest that these two systems initially compete with each other during the acquisition phase, and then become more integrated and interdependent after extended training ^13,14^.

The reinstatement of experience-dependent neural activity during sleep is thought to be a critical element of memory consolidation, allowing new, fragile memory traces to stabilize and reorganize over time ^15^. Supporting this hypothesis, rodent studies showed that learning-related neural patterns of hippocampal place cell activity are spontaneously replayed during sleep, in particular during non-rapid eye movement (NREM) sleep ^16,17^, which does not preclude an important role for REM sleep in memory consolidation (see e.g. ^18,19^). Likewise, human studies evidenced the experience-dependent reactivation of neural activity during post-training sleep, in brain areas previously activated at learning during wakefulness ^4, 20–23^. Regarding spatial memories, a positron emission tomography (PET) study reported increased activity during NREM sleep in navigation-related hippocampal and parahippocampal regions in participants trained to explore a virtual city before sleep, as compared to subjects trained to another, non-hippocampus-dependent procedural memory task ^24^. Moreover, overnight gains in navigation performance were correlated with the amplitude of hippocampal activity during NREM sleep. Besides, functional magnetic resonance imaging (MRI) and magnetoencephalography (MEG) investigations showed that post-learning sleep (as compared to the same time spent awake) putatively leads to the reorganization and/or the optimization of the brain patterns underlying the delayed retrieval of topographical ^5,25^, declarative ^26,27^ or motor procedural ^28–31^ memories. In particular, using a spatial navigation task, it was found that post-learning sleep leads to a restructuration of the neuronal underpinnings of behavior ^5^. Navigation in a recently learned environment, initially subtended by a hippocampus-based spatial strategy, became more contingent on a response-based strategy mediated by the striatum a few days later when participants were allowed to sleep the first night after learning^5,25^, suggesting that post-learning sleep contributed to the automation of the navigation behavior.

While providing us with relevant knowledge regarding the brain networks supporting cognitive functioning during a learning session or at delayed retrieval, task-based fMRI only indirectly explores the offline consolidation processes subtending the maintenance and optimization of novel information outside of the learning or testing environments, as it does not directly measure brain activity during the offline consolidation period. We showed using fMRI that novel topographical learning actually modulates hippocampal responses at wake during a subsequent, unrelated attention task ^32^. Modulated brain activity during post-training wakefulness was suggested to shape and reinforce the functional connections between learning-related cerebral structures and other brain regions, thus contributing to the consolidation process ^32^. Another experimental approach to investigate the offline evolution of the cerebral correlates of recently formed memories without the confounding effect of concurrent task practice is to document the experience-driven changes in intrinsic brain connectivity during a post-training resting state, which can be done using resting state fMRI (rfMRI) ^33^. For instance, Woolley et al. (2015)^34^ found increased functional connectivity between the left posterior hippocampus and the dorsal caudate nucleus in the resting state after (vs. before) navigation in a virtual water maze, which was correlated three days later with offline gains in behavioral performance. Likewise, functional connectivity changes with the hippocampus were investigated after 45 minutes of spatial route learning in a subsequent study ^35^. Results showed that route learning increased synchronization within a network comprising the left hippocampus, the right posterior and anterior inferior temporal gyri and the right intraparietal sulcus. Functional connectivity also increased between the precuneus and the left insula, and between the precuneus and the left hippocampus. These data indicate that rfMRI represents a sensitive approach to investigate the offline processes subtending the consolidation of novel memories.

From an ecological perspective, topographical learning in unfamiliar environments such as unknown cities may usually span over several days or even weeks, the explorer having the opportunity to visit and revisit different but related spatial units, and to integrate the knowledge acquired in previously learned areas with his/her ongoing exploration. In this context, rfMRI provides a valuable tool to better understand the dynamic processes of integration and consolidation following learning or relearning, and their relationships with post-training sleep. In the present study, we investigated rfMRI activity and connectivity changes immediately after spatial navigation learning, at delayed retrieval 4 days later modulated by sleep versus total sleep deprivation on the first post-learning night, and then after a relearning episode in an extended environment encompassing the initial map. To do so, we analyzed resting-state activity using both functional connectivity (that estimates the changes in connectivity between regions) and Amplitude of Low Frequency Fluctuation (ALFF; that estimates spontaneous activity at rest, regardless of the temporal correlation) approaches. Additionally, task (navigation)-based fMRI activity was recorded on Day 1 and Day 4. We hypothesized that sleep-dependent changes in intrinsic functional connectivity in the resting state would parallel and/or complement the previously reported reorganization in task-related brain networks supporting navigation, from hippocampal to striatal and frontal areas^5,25^. Furthermore, we expected further functional connectivity changes between the hippocampus, striatum and frontal areas after renewed spatial learning in the extended version of the initial environment, reflecting ongoing integration with previously consolidated knowledge.

## METHODS

### Participants and General Procedure

In a first step, 51 healthy volunteers gave their written informed consent to participate in a single experimental session (Day 1) of this research approved by the Ethics Committee of the University of Liege, Belgium. One participant was excluded due to abnormal structural MRI scan. The final sample consisted of 50 participants (26 males, 24 females; mean age, 23.5 years; range, 19-32 years). A subset of 34 participants (18 males, 16 females; mean age, 23.5 years; range, 19-31 years) additionally gave consent to further participate in a sleep deprivation vs. regular sleep protocol with a second experimental session (Day 4) (Figure 1). All participants were paid for their participation. Self-reported questionnaires assessed sleep habits (PSQI; Buysse et al., 1989) and circadian topology (MEQ; Horne and Ostberg, 1976) for the past month before inclusion. Participants had to be right-handed (Laterality ^38^ Quotient Score > 70). Participants with extreme circadian chronotype (MEQ score > 69 or < 31),and bad sleep quality (PSQI score > 12), prior history of neurological or psychiatric disorder, or taking psychotropic drugs were not included in the study.

**Figure 1.**
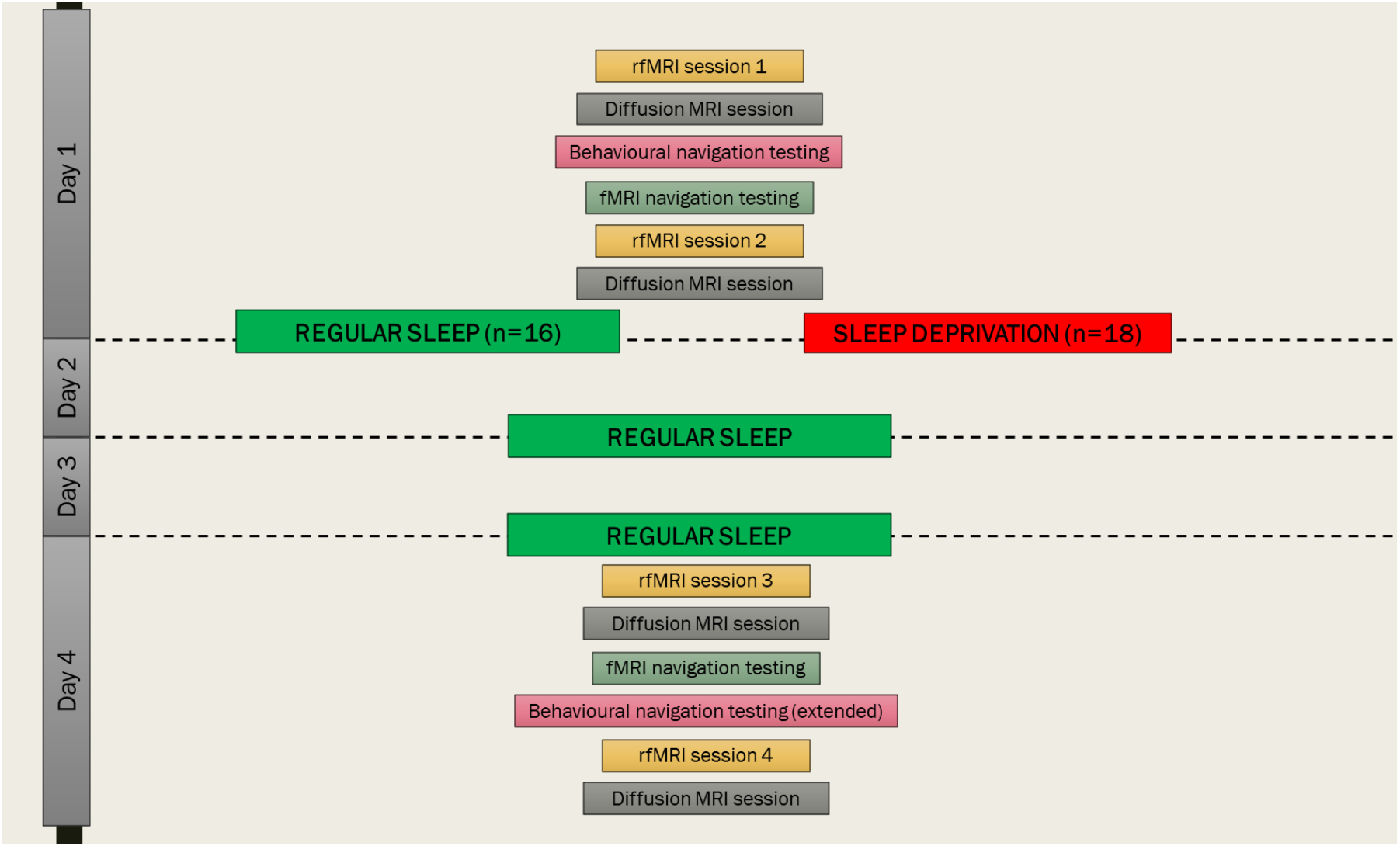
Experimental Design. On Day 1, all participants (n=50) explored the 1st level of the virtual town, and were scanned (5 minutes) in the resting state before (rfMRI 1) and after (rfMRI 2). A subsample of the participants (n=34) was administered immediately afterward(s?) a navigation test while scanned with fMRI (fMRI Test 1; immediate retrieval), and then either slept normally (n = 16) or were sleep deprived (n = 18) during Night 1. All participants slept normally on Nights 2 and 3. At Day 4, they were first scanned in the resting state (rfMRI 3) then were administered again a navigation test under fMRI (fMRI Test 2; delayed retrieval). Afterward, they were trained on an extended version (1st and 2nd levels) of the virtual town, and were scanned a last time in the resting state (rfMRI 4). Finally, participants were tested on the extended version of the virtual town outside of the MRI environment (Retrieval, extended). A diffusion tensor imaging MRI acquisition was performed at each MRI session (data not reported here).

On Day 1, the experimental session was conducted between 1:00 pm and 8:00 pm (Figure 1). Participants (n = 50) were first scanned during a 5-minutes resting state session (rfMRI 1), then had 45 minutes outside of the scanner to explore and learn the first level of the virtual city, followed by a behavioural navigation test where they had to reach as fast as possible 10 targets within the environment (see below). They were then scanned again during a resting state session (rfMRI 2). Afterwards, a subset of participants was enrolled in the sleep deprivation vs. regular sleep protocol (n = 34) and scanned using fMRI while performing again the navigation test (i.e., 10 targets to reach) for approximately 8-minutes (see Figure 1 These participants were randomly assigned either to the regular sleep ([RS], n = 16, 9 males) or to the sleep deprivation ([SD], n = 18, 10 males) condition. Participants in the RS group were allowed normally sleeping at home, and requested to follow regular sleep habits for the 3 following nights. In the SD group, participants were kept awake in the laboratory during the first post-learning night. During this night, participants’ physical activity was maintained as low as possible under constant supervision by the experimenters. They remained most of the time in a seated position, and were allowed to read, chat, play quiet games or watch movies. Participants regularly received isocaloric meals during the night, and water ad libitum. At 8:00 am, they were allowed to leave the lab, and instructed to have non-dangerous daytime activities and to abstain as much as possible from napping until bedtime. They then slept normally at home the two following nights. Sleep quality and quantity for each night from before the learning session (Day1) to the last testing session (Day4) was assessed using a daily standardized questionnaire ^39^ and controlled by visual inspection of actimetric data acquired through continuous actimetric recordings (ActiGraph, wGT3X-BT Monitor, USA).

On Day 4, participants came back to the laboratory for a second experimental session. First, they were scanned in the resting state (rfMRI 3), then scanned using fMRI while engaged in a navigation retrieval test similar to Day 1 (i.e., 10 targets to reach). To control for potential circadian confounds, fMRI scanning took place at the same time of day as the initial testing for each individual. Afterwards, participants were trained outside of the MRI scanner for an additional 40 minutes on an extended version of the navigation task (first and second levels of the city, see below), in which were also incorporated the targets and environments of the initial training. They were then scanned again in the resting state (rfMRI 4), and finally tested on the extended version of the navigation task outside of the MRI environment. A high-resolution structural MRI scan was also acquired for each participant at the end of the brain imaging session, on either Day 1 or Day 4. Additionally, MR diffusion images (duration 12 minutes) were acquired prior to rfMRI 1 and rfMRI 3, as well as after rfMRI 2 and rfMRI 4 (data not reported here).

### Material and Experimental Conditions

Participants were trained in a virtual environment used in prior studies ^5,40^ and adapted from a commercially available computer game (Duke Nukem 3D; 3D Realms Entertainment, Apogee Software, Garland, TX) using the editor provided (Build, Ken Silverman, Realms Entertainment). The environment is a complex town made of two levels communicating through two teleports (Figure 2). Each level is composed of three districts, distinct from each other by a different visual architecture, distinctive objects along the streets, and specific background sounds and music. Each district contains one target location identified by a rotating medallion. Participants navigated the town at ground level using a four-direction keypad with their right hand, at a constant speed of 1.25 virtual unit per second.

**Figure 2.**
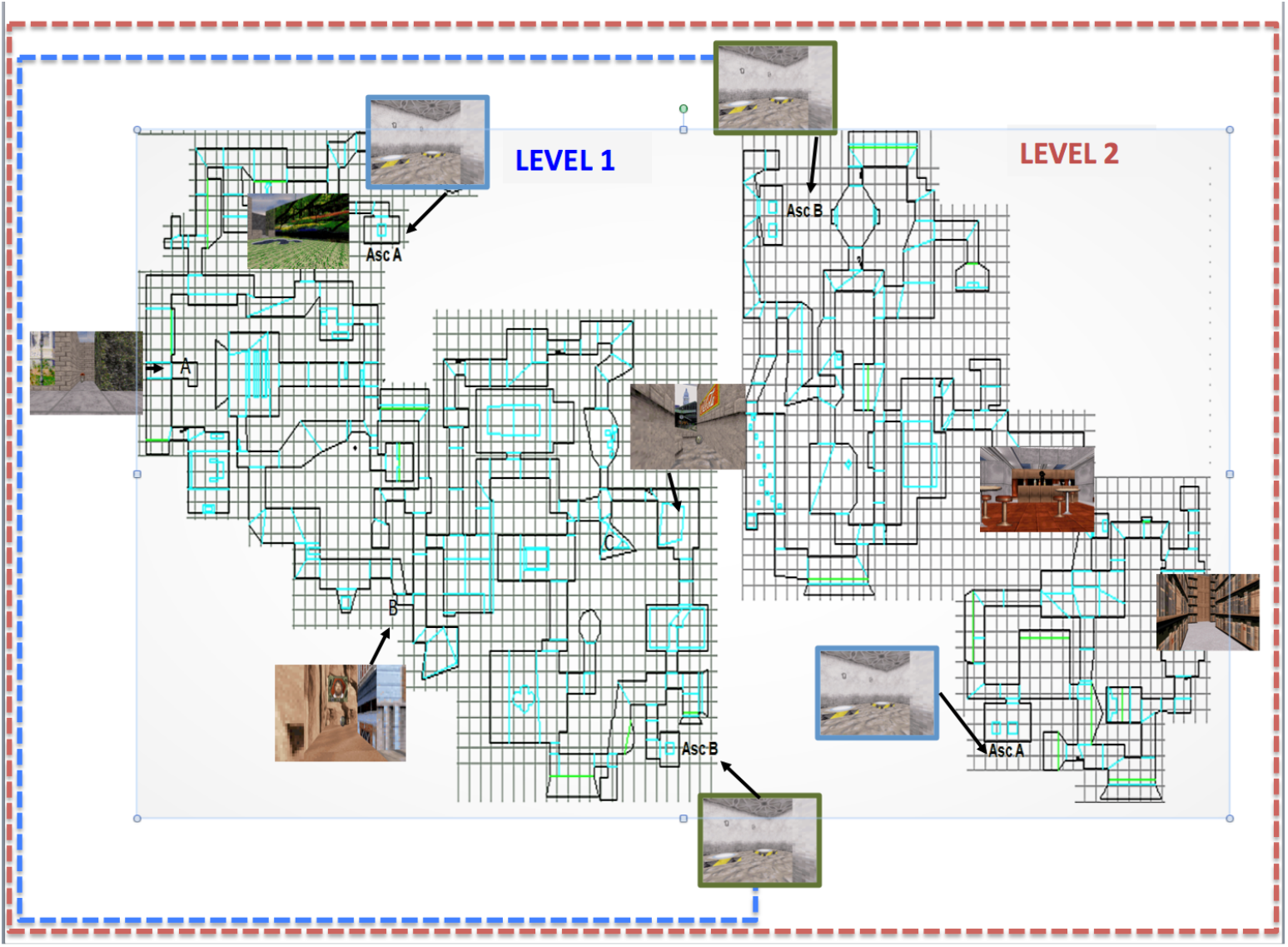
Schematic representation of the map for the immediate and delayed testing sessions in normal condition (starting point on LEVEL 1) and extended (starting point on LEVEL 2), respectively. The targets for both conditions are located on LEVEL 1 (A, B, C) and the two levels are connected through elevators (AscA and AscB).

#### Behavioural sessions

During behavioural sessions outside of the scanner (Learning [Day 1] and Relearning [Day 4] phases), the virtual environment was presented on a personal computer laptop (screen size, 17 inches), and participants were trained for about 45 minutes.

On Day 1 (Learning), the training consisted of four 8-minutes blocks of free exploration within the first level of the virtual environment, each block starting from one of the three possible targets. Participants were explicitly instructed to learn the spatial layout of the streets, districts, and target locations by moving freely within the environment. During the entire training session, pictures of the three target locations and their associated names were continuously available. At the end of each 8-minute block, participants were exposed to a test trial similar to the one proposed in the fMRI environment, i.e. they were assigned a starting point and instructed to reach a given target location as fast as possible and in a maximal time of 28 seconds (i.e., the shortest possible duration from starting point to target destination). At the end of the four blocks of free exploration, participants were exposed to 10 test trials in a row, to be performed later in the fMRI setting (see below).

At Day 4 (Relearning), participants were trained again for four 8-minutes blocks of free exploration within an environment composed of the initial three districts (Level 1) and of another level (Level 2) made of three new districts. They were informed that both levels were connected by two teleports positioned at opposite locations, and instructed to learn the layout of Level 2 as well as the locations of each teleport, in order to be able to reach the initial targets located on Level 1, starting from random locations in Level 2. Each block started at one of the two teleports. At the end of each 8-min training block, participants performed a test trial, i.e. they were assigned a starting point in Level 2 and asked to reach, as fast as possible, one of the initial targets in Level 1 within a 60-second window (i.e., the shortest possible duration from Level 2 starting point to Level 1 target destination). After training and [rfMRI 4] acquisitions, participants were administered a final test of ten 60-sec trials outside of the MRI environment.

#### Task-based fMRI

During task-based fMRI (Immediate retrieval fMRI Test 1 and Delayed retrieval fMRI Test 2; Figure 1), volunteers performed a series of navigation tests in the virtual town. For each test, they were assigned a starting point and instructed to reach, as fast as possible, a given target location within the time limit of 28 seconds. Because the duration of the shortest path between the starting point and the target is 28 s at constant speed, reaching the destination could only be achieved if participants selected the ideal path and did not stop at all during the navigation. The fMRI test session consisted of 10 test blocks, each lasting for 28 s and alternating with a pause menu displayed for a variable duration ranging 10–17 s.

For both Day 1 and Day 4 behavioural and fMRI testing sessions, participants’ trajectories were analysed a posteriori based on video recordings, and a quantitative measure of performance was computed at each block as the shortest distance remaining between the subject’s actual location at the end of the time limit for the test trial and the location of the target destination ^5,25^.

#### Resting state rfMRI

For rfMRI sessions 1, 2, 3 and 4,_participants were instructed to relax, keep their eyes open and fixate for 5 minutes the cross displayed on the screen.

### Brain imaging data acquisition

Brain MRI data were acquired on a whole-body 3T scanner (Magnetom Prisma, Siemens Medical Solutions, Erlangen, Germany) operated with a 20- or 64-channel receiver head coil.

For both task-based and resting-state fMRI, multi-slice T2*-weighted functional images were acquired with a gradient-echo echo-planar imaging (EPI) sequence using axial slice orientation and covering the whole brain (36 slices, FoV = 216×216 mm^2^, voxel size 3×3×3 mm^3^, 25% interslice gap, matrix size = 72×72×36, repetition time (TR) = 2260 ms, echo time (TE) = 30 ms, flip angle (FA) = 90 deg).

For task-based fMRI Retrieval Test 1 and Test 2 sessions, 210 functional images were recorded with the 20-channel head coil. The virtual environment was displayed on a screen positioned at the rear of the scanner that the subject could comfortably see through a mirror mounted on the head coil. Navigation was done using 4 buttons on a commercially available MRI compatible keypad system (fORP; Current Design, Vancouver).

For each resting state session (rfMRI 1-4), 132 functional images were obtained with the 64-channel head coil.

For anatomical reference, a high-resolution T1-weighted structural image was acquired for each subject (T1-weighted 3D magnetization-prepared rapid gradient echo (MPRAGE) sequence, TR = 1900 ms, TE = 2.19 ms, FA = 9 deg, inversion time (TI) = 900 ms, FoV = 256×240 mm^2^, matrix size = 256×240×224, voxel size = 1×1×1 mm^3^, acceleration factor in phase-encoding direction R=2, 64-channel head coil).

### Task-based fMRI Data Analysis

Preprocessing and analysis of functional volumes were performed using the Statistical Parametric Mapping software SPM12 (Wellcome Department of Cognitive Neurology, London) implemented in MATLAB R2012B (Mathworks, Sherbom, MA). The five initial volumes of each time series were discarded to avoid T1 saturation effects. Preprocessing of individual data included realignment (2-step realignment on the first volume of the series), correction for geometric distortions caused by magnetic fields based on the Field Map Toolbox 41, co-registration of functional and anatomical data, spatial normalization into standard stereotactic Montreal Neurological Institute (MNI) space, and spatial smoothing using a Gaussian kernel of 6-mm full width at half maximum (FWHM). Excessive head movements of more than 4 mm of translation or 4 degrees of rotation in any direction were considered exclusion criteria. No participant exhibited excessive head motion.

Functional data were analysed using a mixed-effects model aimed at showing stereotypical effect in the population from which the subjects are drawn ^42^. For each subject, a first-level intra-individual analysis tested effects of interest by linear contrasts convolved with a canonical hemodynamic response function, generating statistical parametric maps. Movement parameters derived from the realignment phase were included as confounding factors. Cut-off period for high-pass filtering was 128 s due to the block design alternating 28-seconds navigation blocks with 10-17 seconds breaks. Additionally, navigation performance was added to the model to test possible relationships between neuronal activity in navigation–related areas and behavioural performance. Navigation-related regional BOLD response modulated by navigation performance was computed at the within-subject level. Since no inference was made at this fixed effects level of analysis, individual summary statistic images were thresholded at p<0.95 uncorrected, then further spatially smoothed (6 mm FWHM Gaussian kernel). Individual summary statistics images were then entered in a second-level analysis, corresponding to a random effects (RFX) model, to evaluate commonalities and differences in brain response between the RS and SD groups, and between immediate and delayed Retrieval Test sessions. The resulting set of voxel values for each contrast constituted a map of the *t* statistic [SPM(T)]. Statistical inferences were obtained after correction for multiple comparisons at the voxel level (Family Wise Error (FWE) correction) (p < 0.05) in the whole brain, or after small volume correction (SVC) in five regions of interests (ROIs), selected based on published reports showing their involvement in topographical/spatial learning (Boccia et al., 2014; Brown et al., 2014; Epstein et al., 2017; Reagh and Yassa, 2014; Robin et al., 2015), i.e. bilaterally the hippocampus, the parahippocampal gyrus, the retrosplenial and medial parietal cortices (Brodmann areas 29, 30 and 31), the dorsal striatum (caudate and putamen) and the entorhinal cortex. Positions and dimensions of ROI were obtained with the Automated Anatomical Labelling (AAL) atlas^46^.

### Resting state data Analysis

#### Preprocessing

Preprocessing of rfMRI data was carried out using the SPM12 software and the Data Processing Assistant for Resting-State fMRI (DPARSF) ^47^. The first 10 volumes of each time series were discarded to avoid T1 saturation effects. Functional images were realigned to the first volume to correct for head motion. Participants with translation superior to 2.0 mm in any direction, or angular motion superior to 2.0 along any axis were excluded from the analysis. Following this criterion, one participant was excluded. The final samples for rfMRI analyses therefore consisted of 49 participants for day 1, and 34 participants for day 4. Subsequently, each individual structural image (T1-weighted MPRAGE image) was co-registered to the mean functional image after motion correction using a linear transformation. The transformed structural images were then segmented into gray matter (GM), white matter, and cerebrospinal fluid using Diffeormorphic Anatomical Registration Through Exponentiated Lie Algebra (DARTEL). To further reduce potential confounds of head motion, a Friston-24 correction ^48^ was applied based on six rigid body head motion parameters. Signals from cerebrospinal fluid and white matter were also entered as confound covariates to reduce possible effects of physiological artifacts. The corrected functional volumes were spatially normalized to the Montreal Neurological Institute (MNI) space, and re-sampled to 3-mm isotropic voxels using DARTEL. Resultant normalized functional images were de-trended and spatially smoothed with an 8-mm full-width-at-half-maximum (FWHM) Gaussian kernel.

#### ALFF analysis

Smoothed images were band-pass filtered (0.1–0.01 Hz) to reduce low-frequency drifts and high-frequency noise before the Amplitude of Low Frequency Fluctuations (ALFF) analysis aimed at investigating local changes in resting state activity. ALFF individual maps were computed with DPARSF and entered into second-level analyses to examine increases in ALFF signal associated with a spatial learning episode on Day 1 and Day 4, and to investigate if functional changes associated with learning in the extended version of the environment differed between groups on Day 4. ALFF changes associated with performance were also investigated by entering behavioral performance as a covariate of interest at the second level (RFX) of the analysis. Statistical inferences were computed at the whole brain level based on a two-tailed permutation test with 5000 permutations and a threshold-free cluster enhancement (TFCE) correction within a grey matter mask (obtained by thresholding an a priori grey matter probability map in SPM12; threshold = 0.25). Results were considered significant at p<.05 after a false discovery rate (FDR) correction for multiple comparisons. Region of interest analyses were also conducted within the five a priori ROIs defined above based on the literature ^8,11,43–45^. Clusters in which activations were found to correlate with performance in the task-based fMRI analysis on Day 1 and/or Day 4 were also examined as regions of interests for Day 1 (3 clusters) and Day 4 (0 clusters) ALFF analyses.

#### Functional Connectivity analysis

Functional connectivity was analyzed with seed-voxel correlation mapping using the CONN-fMRI toolbox 13.i for SPM (http://www.nitrc.org/projects/conn)^49^. Preprocessed data (see “Preprocessing section”) were temporally filtered using a band-pass filter to retain frequencies between 0.008 to 0.09 Hz. Noise signals, such as cerebrospinal fluid, white matter, movement parameters and time-series predictors of global signal were further removed from the images using the component-based noise correction method (CompCor) ^50^. Averaged signals were extracted from 10 different seeds, defined using the Automated Anatomical Labeling (AAL) atlas: left and right hippocampus, parahippocampus, dorsal striatum, retrosplenial-medio parietal cortex and entorhinal cortex (regions of interest taken from ^8,11,43–45^). Clusters found to correlate with performance in the task-based fMRI and ALFF analyses on Day 1 and Day 4 were also used as seeds for Day 1 and Day 4 analyses respectively, yielding 7 extra seeds for Day 1 and 2 extra seeds for Day 4. Temporal correlations of resting-state BOLD signal time series between these seeds and the rest of the brain were examined using a General Linear Model approach. Individual connectivity maps were computed for each seed and entered into a second level analysis to examine changes in connectivity following navigation learning on Day 1, and to investigate if functional changes associated with learning in the extended version of the environment differed between groups on Day 4. Behavioral performance following learning was used as a covariate of interest (or as a weighting factor of BOLD signal). Statistical inference for each seed and contrast were first based on a family-wise error correction (p < .005) at the cluster-level with an uncorrected threshold of p < .001 at the voxel level. A Bonferroni correction was then applied based on the number of seeds for Day 1 (n=17) and Day 4 (n=12) analyses separately. Results were thus considered significant at a threshold of p = .0014 (0.05/17) for Day 1 analyses and p = .002 (0.05/12) for Day 4 analyses.

The data that support the findings of this study are available from the corresponding authors, MD and PP, upon reasonable request.

## RESULTS

### Behavioral results

#### Sleep data

Sleep duration and quality were estimated by means of self-reports of the nights preceding Day 1 and Day 4. Mean sleep duration was not significantly different between groups on the night before Day 1 (RS = 437 ± 60 min, TSD = 457 ± 46 min; p > .2) and the night before Day 4 (RS = 436 ± 55 min, TSD = 459 ± 48 min; p > .2). Within each group, duration of each night did not differ significantly (RS group: p > .8; TSD group: p > .7). Likewise, a similar analysis conducted on subjective sleep quality, assessed by means of a 6-point scale, failed to evidence any between-group differences. Subjective sleep quality was equivalent between groups for each night (night before Day 1: RS = 3.7 ± 1.2, TSD = 3.4 ± 1.1; p > .5; night before Day 4: RS = 3.7 ± .7, TSD = 3.7 ± .9; p > .9). Finally, within group comparisons did not reveal any significant difference in sleep quality between both nights (RS group: p > .9; TSD group: p > .1).

#### Navigation Performance

As a reminder, for each of the 10 place finding tests administered during fMRI sessions at Day1 (Immediate retrieval) and Day4 (Delayed retrieval), participants were instructed to reach, as fast as possible, a given target from one designated starting point within 28 seconds. Distance remaining to reach destination was the estimate of navigation performance (see *Methods*). Mean distance scores per session for the SD group were 17.17 (arbitrary units, SD 8.29) and 17.53 (SD 11.1) at immediate (Day 1) and delayed retrieval (Day 4), respectively. Distances scores for the RS group were 15.12 (SD 13.15) at immediate and 15.47 (SD 12.2) at delayed retrieval, respectively. A two-way ANOVA computed on within-session individual mean scores with group (SD vs. RS) and retrieval session (Immediate vs. Delayed) factors did not reveal any significant main or interaction effect (all ps > 0.1). Likewise, at testing after the second learning session on the extended version of the town, mean performance did not differ statistically (t = − .2; p = .8) between the RS (arbitrary units, 15.8, SD 19.10) and SD (17.3, SD 19.7) conditions.

These results suggest that (1) both groups gained knowledge of the virtual town that persisted 3 days after learning, that (2) sleep deprivation on the first post-training night did not alter subjects’ ability to find their way in the town at delayed retrieval (which does not preclude the use of qualitatively different strategies for a similar performance), and that (3) learning novel routes and integration with prior knowledge in the extended version of the town was similar in the RS and SD conditions.

### Task-based FMRI

#### Navigation-Related Brain Activity

A conjunction analysis disclosed increased blood–oxygen level-dependent (BOLD) responses in an extended hippocampo-neocortical network during place finding both at immediate (Day 1) and delayed (Day 4) retrieval sessions. At immediate retrieval, navigation-related activity was mostly found in the right hippocampus (32; −40; −4 mm in standard stereotactic space; Z = *4.55, k = 30*; 26, −34, 48 mm; Z = 3.84; k = 12, p_svc_ = .04) and surrounding structures as well as in occipital, parietal, frontal and cerebellar areas (Supplementary Table 1). At delayed retrieval, increased BOLD activity was evidenced in the same set of brain structures, except for the right hippocampus in which activation did not survive correction for multiple comparisons (Supplementary Table 2). Correlation analyses disclosed a positive relationship at immediate retrieval between individual performance score (i.e., reversed score based on the distance toward destination, higher scores meaning better performance) and navigation-related responses in the left (−22; 0; −10, Z = 4.14, k=74 p_svc_ = 0.027) putamen. At delayed retrieval, no significant correlation was found between performance and navigation-related activity, either at the whole brain level or within ROIs.

#### Post-training Sleep-Dependent Reorganization of Brain Activity during Navigation

A group (SD vs. RS) by session (Day 1 vs. Day 4) ANOVA conducted on navigation-related brain activity did not disclose any main or interaction effect, either at the whole brain level or within ROIs. A similar analysis conducted on navigation-related regional BOLD responses modulated by navigation performance also did not yield significant results.

### Resting state ALFF

#### Spatial Learning-Related Changes

On Day 1, local ALFF increased following learning (rfMRI 2 vs. rfMRI 1) in a parietal medial cluster encompassing parts of the precuneus and of the posterior cingulate gyrus (6; −45; 21 mm; intensity = 479.36; k = 268, p_FDR_<0.05) and an extended cluster encompassing frontal and parietal regions (6; −30; 60; intensity = 513.80; k = 409, p_FDR_<0.05). On Day 4, local ALFF increased following learning in a parietal cluster with the maximal peak in the left precuneus (−3; −57; 15; intensity = 788.80; k = 4072, p_FDR_<0.05), a cerebellar cluster (−36; −60; −42; intensity = 528.01; k = 130, p_FDR_<0.05), and two clusters located in the left rectus (0; 24; −21; intensity = 623.98; k = 271, p_FDR_<0.05) and the right lingual gyrus (: 15; −99; −12; intensity= 593.63; k = 463, p_FDR_<0.05).

On Day 1, there was a correlation between navigation performance and increased ALFF in parietal medial (3; −51; 27; intensity 399.51; k = 74, p_FDR_<0.05), right parahippocampal gyrus (33; −18; −30; intensity 50.38; k = 24, p_svc_ <0.05) and left posterior cingulate (−3; −54; 27; intensity = 104.69; k = 57, p_svc_ <0.05) clusters. On Day 4, navigation performance correlated with local ALFF in the left hippocampus (−15; −6; −15; intensity = 72.10; k = 24, p_svc_ <0.05), and the left parahippocampal gyrus (−24; −6; −33; intensity = 95.99; k = 61, p_svc_ <0.05) clusters.

#### Post-training Sleep-Dependent Learning-Related Changes

Between-group comparison revealed no significant difference in ALFF changes associated with learning on Day 4 when comparing RS and SD groups.

### Resting state FC

#### Spatial learning-related changes

As a reminder, seeds for connectivity analyses were defined based on *a priori* ROIs (10 seeds) and on clusters found to correlate with performance in the task-based fMRI or ALFF analyses (see Methods section), for a total of 16 seeds for Day 1 and 12 seeds for Day 4. All analyses are Bonferroni corrected for multiple comparisons. On Day 1 (corrected significance threshold p < .05/16 = .0031), increased functional connectivity (from pre-learning [rfMRI 1] to post-learning [rfMRI 2] resting state session) was found between the left hippocampus and the right middle frontal gyrus (32; 4; 56 mm, k = 247, T=4.95, cluster p_FWE_= 0.0023), between the right entorhinal cortex and three clusters located in the right temporal (52; −18; −24, k = 351, T=5.46, cluster p_FWE_= 0.000001) and left temporal (−58; −16; −24, k = 401, T=5.02, cluster p_FWE_= 0.000066) cortices, and in the right posterior cingulum (10; −42; 28, k = 605, T=4.18, cluster p_FWE_= 0.000197), as well as between the left entorhinal cortex and a cluster located in the cerebellum (−30; −40; −34, k = 310, T=4.88, cluster p_FWE_= 0.000497). Based on seeds taken from task-based correlation analyses between navigation-related activity and performance, spatial learning on Day 1 was associated with increased connectivity between the left parahippocampal gyrus and the right precuneus (6; −60; 44, k = 330, T=5.09, cluster p_FWE_= 0.000211), the right precuneus and the right middle frontal gyrus (30; 4; 38, k = 324, T=4.26, cluster p_FWE_= 0.000343) and the left posterior cingulate cortex and the right supramarginal gyrus (58; −34; 48, k = 232, T=4.57, cluster p_FWE_= 0.000377; see Table 1 for detailed results). No change in brain functional connectivity correlated with performance.

**Table 1:** Spatial learning related changes in resting state functional connectivity on Day 1 (rfMRI2>rFMRI1, all subjects considered). MNI coordinates (mm) indicate peak-voxel location. FWE= Family Wise Error. *= seeds obtained from task-related fMRI or ALFF analysis. K= cluster size expressed in number of voxels.

| Seed | Target brain region | x y z | K | T | Cluster p-FWE (threshold < .0029) |
| --- | --- | --- | --- | --- | --- |
| Left Parahippocampal Gyrus* | Right Precuneus | 6 -60 44 | 330 | 5.09 | 0.000211 |
| Right Precuneus* | Right Middle Frontal Gyrus | 30 4 38 | 324 | 4.26 | 0.000343 |
| Cingulum_Post_L* | Right Supramarginal Gyrus | 58 -34 48 | 232 | 4.57 | 0.000377 |
| Left Hippocampus | Right Middle Frontal Gyrus | 32 4 56 | 247 | 4.95 | 0.002300 |
| Right Entorhinal Cortex | Right Inferiotemporal Gyrus | 52 -18 -24 | 351 | 5.46 | 0.000001 |
|  | Left Middle Temporal Gyrus | -58 -16 -24 | 401 | 5.02 | 0.000066 |
|  | Right Post Cingulate Gyrus | 10 -42 28 | 605 | 4.18 | 0.000197 |
| Left Entorhinal Cortex | Left lobule VI of cerebellar hemisphere | -30 -40 -34 | 310 | 4.88 | 0.000497 |

On Day 4 (corrected significance threshold p < .05/12 = .0041), learning in the extended version of the task triggered increased functional connectivity (from the pre-learning [rfMRI 3] to the post-learning [rfMRI 4] resting state session) between the left hippocampus and the right postcentral gyrus (36; −30; 70, k = 249, T=4.34, cluster p_FWE_= 0.002026). This functional connectivity change did not correlate with performance.

#### Sleep-dependent changes in resting state functional connectivity

At Day 4 during the first resting state session (rfMRI 3) before extended learning in the maze, RS participants exhibited higher functional connectivity than SD participants between the right dorsal striatum and the left middle frontal gyrus (−34; 30; 42, k = 272, T=4.71, cluster p_FWE_= 0.001851) (Figure 3).

**Figure 3.**
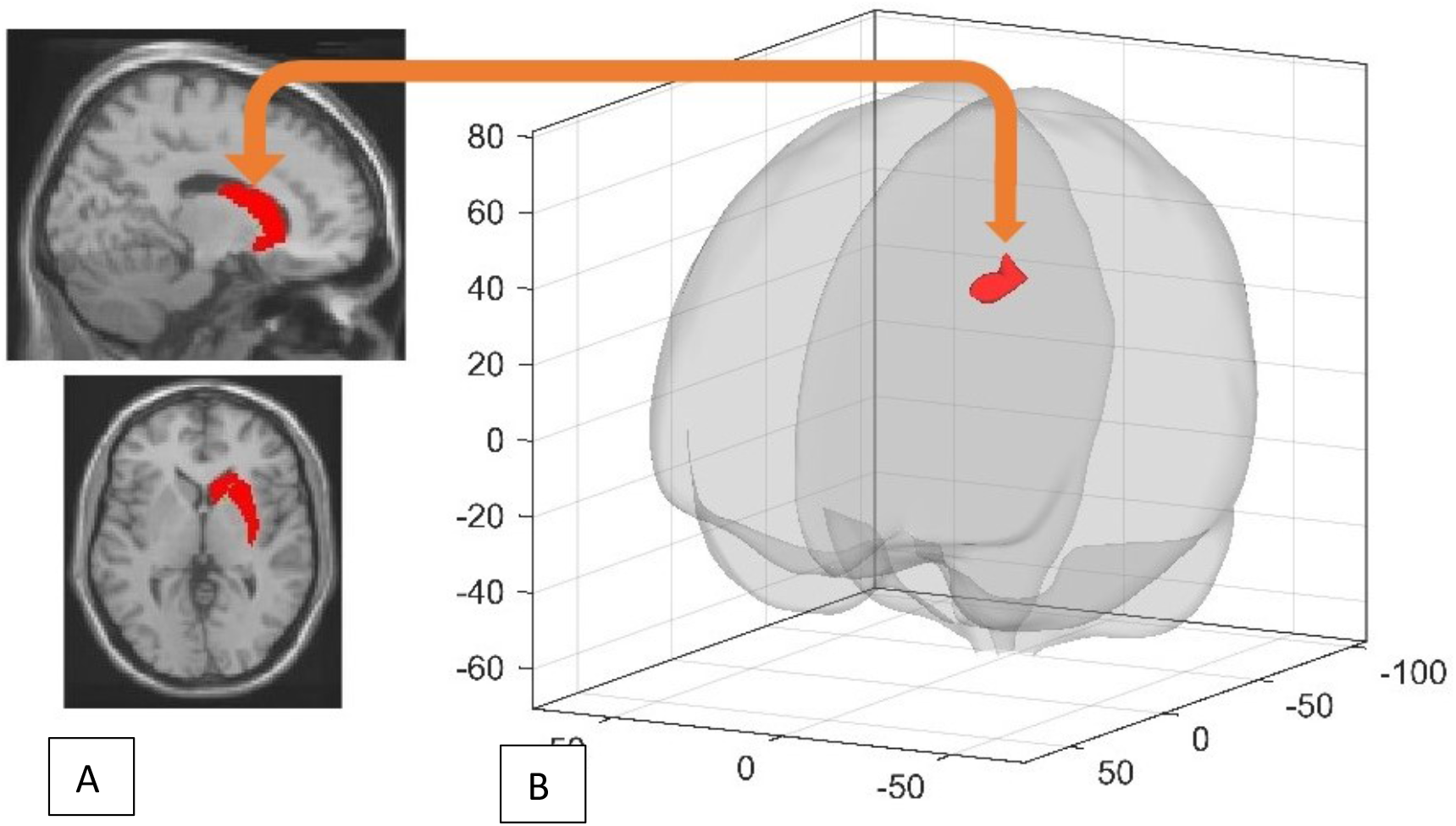
RS group exhibited higher FC between right dorsal striatum seed (A) and middle frontal gyrus target cluster (B) (cluster visualised with “glass display” available in CONN toolbox).

#### Post-training sleep-dependent learning-related changes in resting state functional connectivity

A Group (RS vs. SD) by resting state Session (pre-extended learning [rfMRI 3] vs. post-extended learning [rfMRI 4]) interaction analysis investigated sleep-dependent changes in resting state functional connectivity after learning on the extended version of the town at Day 4. This analysis disclosed increased functional connectivity from pre- to post-learning in the SD as compared to the RS condition, between the left retrosplenial cortex and the left superior frontal gyrus (−16; 12; 48, k = 738, T=5.91, cluster p_FWE_ < 0.000001), between the left retrosplenial cortex and the right middle frontal gyrus (28; 6; 56, k = 390, T=5.78, cluster p_FWE_= 0.000047), and between the right entorhinal cortex and the right superior frontal gyrus (18; 24; 56, k = 228, T=5.25, cluster p_FWE_= 0.001829; Table 2). A post-hoc analysis ran in CONN fMRI to investigate the directionality of the interaction effect revealed significant decreases in FC in the RS group vs. increases in the SD group (Left retrosplenial cortex and left superior frontal gyrus: RS, β_pre-learning_ = 0.28; β_post-learning_ = 0.05 ,T=4.44, p<0.001, SD, β_pre-learning_ = 0.12; β_post-learning_ = 0.26 , T=3.13, p<0.001; left retrosplenial cortex and right middle frontal gyrus: RS, β_pre-learning_= 0.13; β_post-learning_ = −0.07 ,T=4.38, p<0.001, SD, β_pre-learning_ = −0.08; β_post-learning_ = , T=2.94, p=0.004; right entorhinal cortex and right frontal superior gyrus: RS, β_pre-learning_ = 0.07; β_post-learning_ = −0.09 ,T=5.19, p<0.001, SD, β_pre-learning_ = −0.15; β_post-learning_ = − 0.02 , T=2.80, p=0.006) (Figure 4).

**Table 2:** Post training Sleep-Dependent Learning-Related Changes in resting state functional connectivity (rfMRI4>rfMRI3, SD>RS). MNI coordinates indicate peak-voxels location. FWE= Family Wise Error. K= cluster size expressed in number of voxels.

| Seed | Target brain region | x y z | k | T | Cluster p-FWE |
| --- | --- | --- | --- | --- | --- |
| Left Retrosplenial Cortex | Left Superior Frontal Gyrus | -16 12 48 | 738 | 5.91 | 0.000000 |
|  | Right Middle Frontal Gyrus | 28 6 56 | 390 | 5.78 | 0.000047 |
| Right Entorhinal Cortex | Right Superior Frontal Gyrus | 18 24 56 | 228 | 5.25 | 0.001829 |

**Figure 4.**
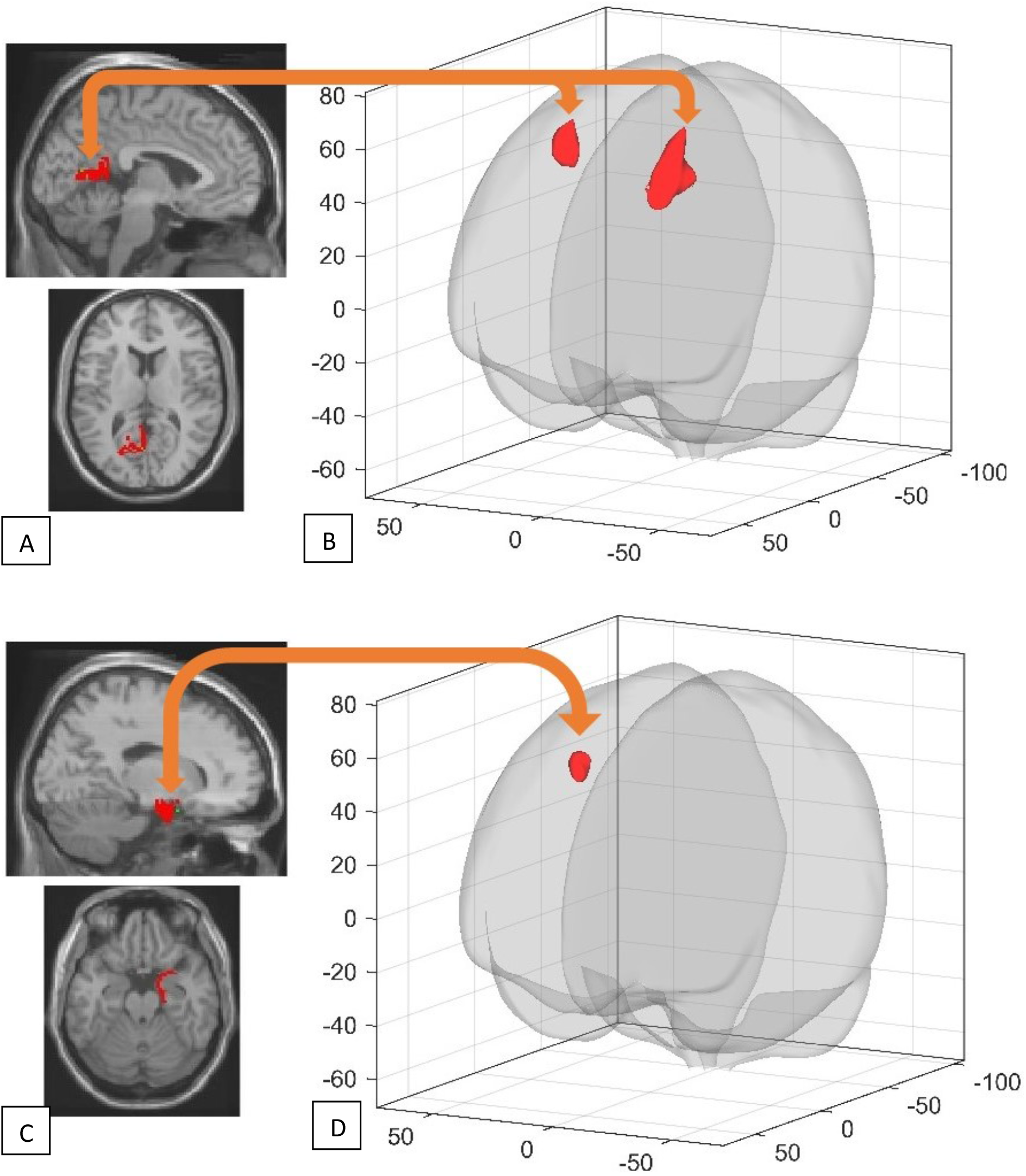
SD group exhibited higher functional connectivity between left retrosplenial cortex seed (A) and two frontal target clusters (left superior frontal gyrus and right middle frontal gyrus)(B), as well as an increased connectivity between right entorhinal cortex seed (C) and a right frontal gyrus target cluster (D) (target clusters visualised with “glass display” available in CONN toolbox).

## DISCUSSION

In the present study, resting-state fMRI was used to investigate the offline changes in brain functional connectivity and intrinsic regional activity potentially associated with learning and relearning following sleep in a spatial navigation task.

In line with previous studies, behavioural performance was similar between groups sleep deprived or sleep rested post-training, not only at immediate but also at delayed retrieval ^5,25^, or when tested after learning the extended version of the task. It suggests that one night of sleep deprivation following training is insufficient to induce observable changes at the behavioural level (see also ^51^ for instance), which does not exclude the development of distinctive neural strategies subtending successful navigation ^5,12,25^. Indeed, prior studies found a shift between neural structures subtending quantitatively equivalent retrieval performance after sleep on the post-training night, from hippocampus to striatum in the virtual navigation task after a few days ^5,25^, and from hippocampus to neocortical frontal regions in other declarative memory tasks after a sleep episode in children ^27^, and after weeks to months in adults ^26,52^.

Learning to navigate in the virtual environment on Day 1 was associated with increased functional connectivity between the left hippocampus and the right middle frontal gyrus, as well as in a network comprising the right entorhinal cortex, bilateral temporal regions and the right posterior cingulum cortex. The right middle frontal gyrus is active during virtual maze exploration ^53^, and important for episodic memory retrieval ^54^, while the cingulum contributes to the sense of self-location ^55^, and the entorhinal cortex plays a key role for the encoding of map-like spatial codes ^56^. Consequently, increased functional connectivity may subtend the integration of egocentric and allocentric spatial representations. When considering as seeds the clusters obtained from task-related fMRI and ALFF analyses, increased functional connectivity was found between the left parahippocampal gyrus and the right middle frontal gyrus, a set of regions centrally related to spatial memory^57–59^. On day 4, learning in the extended version of the task triggered an increase in functional connectivity between the left hippocampus and the right postcentral gyrus, two regions subtending spatial navigation in recently learned environments ^60^. Noticeably, recent reports proposed that besides being the somatosensory area, the postcentral gyrus plays also a role as a high-order navigational system ^61^. In rats, spatial cells in this system elicit similar firing characteristics than the ones in the hippocampus ^61^. To the best of our knowledge, this finding has not yet been replicated in human subjects. Notwithstanding, task-dependent increased functional connectivity between the hippocampus and somatosensory areas may be seen as evidence supporting the hypothesis of a cooperation between these two navigational systems.

The first analysis addressing sleep deprivation effects on resting-state activity was performed as soon as the participants arrived at the laboratory (Day4), before any kind of learning related stimulus was presented. Interestingly, the RS group presented a higher functional connectivity between the right dorsal striatum and and the left middle frontal gyrus, compared to the SD group. Even if there is no concrete evidence that this increase in functional connectivity is related to the task learned on the first day, this results can be still seen as an indication of sleep-deprivation dependent reorganization of brain function in areas that are recruited during learning and navigation.

Learning to navigate in an extended version of the initial environment was associated with increased functional connectivity between navigation-related brain structures (left retrosplenial cortex and right entorhinal cortex) and frontal regions (left and right superior frontal gyrus, right middle frontal gyrus) in participants deprived of sleep the night after initial learning (SD) as compared to those having slept normally (RS). The entorhinal and the frontal cortices are respectively thought to support landmark recognition and strategy planning ^62^ in the context of spatial navigation, while the retrosplenial cortex plays a central role in spatial strategy switching ^63,64^. Stronger functional connections with frontal regions in SD participants might reflect the compensatory brain activity needed to develop different but quantitatively equivalent (thus similar behavioural performance) spatial representation strategies in the context of extended learning in a less strongly consolidated environment. This tentative interpretation should be tested in further studies.

We also investigated changes in the amplitude of low frequencies fluctuation (ALFF) as potential markers of learning and memory consolidation processes ^65,66^. Both in the first (Day 1) and in the second (Day 4) contrast, changes in local brain activity (ALFF) at rest partially overlapped with clusters found active when performing on the navigation task, suggesting the offline continuation of task-related activity during post-training wakefulness^32^. Amongst the regions found active in both online task and offline post-training conditions in the present study, the precuneus was previously involved in self-related mental representations both during rest and spatial information processing ^67^. Post-training activations in the precuneus may thus support the creation of the egocentric spatial map needed for successful topographical navigation. Increased spontaneous activity (ALFF) was also detected at Day 1 in the posterior cingulum, a region that plays an important role in visual-spatial memory due to its connections with hippocampal and parahippocampal regions ^68^. At Day 4, ALFF increased in the right lingual gyrus, a part of the so-called “parahippocampal place area” (PPA) that responds preferentially to complex visual scenes such as landscapes or cityscapes^69^. However, the lack of specific literature relating ALFF with memory consolidation processes calls for caution in the interpretation, and it is premature at this stage to link ALFF changes to the replay of memory traces. Both at Day 1 and Day 4, increased resting state activation in the medial temporal lobe positively correlated with behavioural navigation performance, suggesting that prior learning modulates the activation of this brain region that to some extent reflects successful learning In line with behavioural and task-fMRI results, the group (SD vs. RS) by session interaction on spontaneous activity was non-significant, suggesting that ALFF measurements mostly reflect immediate post-training memory processes at wake, but were not modulated by SD after the first post-training night.

At both immediate and delayed retrieval, task-based fMRI analyses evidenced navigation-related brain activity in a large network including frontal and occipital areas, similar to the ones previously found to support navigation in this task ^5,25^. However, at variance with previous studies from our group reporting significant correlations between performance and medial temporal activity ^5,40^ at immediate retrieval, performance at Day 1 already correlated with striatal activation in the current study. Consequently, we could not replicate the observation of a shift from hippocampal to caudate-dependent spatial strategy in the RS as compared to the SD group ^5,40^, as striatal activation was already present at Day 1. We propose here three possible explanations for this discrepancy with prior reports. First, it should be taken into account that participants spent approximately 20 minutes lying in the MRI machine (for resting-state fMRI and diffusion imaging acquisitions) prior to the training session, which might have augmented stress levels and sleepiness. In animals, these elements were proposed to lead to an increased use of the striatal spatial memory system ^70^. Second, at variance with our prior study ^5^, the behavioral training proposed to participants on Day 1 included at the end a repetition of the 10 tests administered during the subsequent task-based fMRI session. This final test was added to reinforce the memory traces and habituate our participants to the conditions in which they would have to perform next in the scanner. However, as participants experienced already once the navigation in these conditions, this may have eventually resulted (and much faster than we expected) in a form of automation of the navigation, with a shift toward a striatum-based strategy. Third, the results obtained in our prior study now date back more than 12 years. Since that time, the use of geolocation software to navigate in the real world has become a given for the new generations, and especially for our young adult volunteers. Thus, it cannot be excluded that the spontaneous neuronal underpinnings of spatial navigation performance have evolved in unexpected ways due to the quasi-constant availability of external navigation systems that to some extent replace the need to create topographical representations, a hypothesis that needs experimental confirmation. Finally, the study protocol (as the previous ones ^5,25^), does not include a “control condition” where participants train on an exploration experience not implicating learning, which would have given us a more specific aspects of learning dynamics. Similarly, the results regarding the interaction between post-learning sleep deprivation *and* extension of a virtual environment would require further confirmation in a larger cohort by using a control condition in which half of the participant of each group (RS and SD) is trained on a completely unrelated environment on day 4.

Altogether, our results highlight the continuation of navigation-related activity in the subsequent resting state, as evidenced by changes in functional connectivity and the amplitude of low frequencies fluctuation in task-related neural networks. Furthermore, sleep deprivation on the post-training night was associated with increased functional connectivity between navigation-related brain structures when faced to the task to learn a novel but related environment (i.e., the extended version of the city), suggesting the need to recruit more resources to link novel information with possibly less efficiently consolidated existing memory traces.

## Acknowledgements

This study was financially supported by grants from the Fonds de la Recherche Scientifique (FRSM project #7020836, Brussels, Belgium) and from the Excellence Of Science (EOS) FNRS-FWO (MEMODYN project 30446199). At the time of the study, TV was funded by an Université Libre de Bruxelles (ULB) Individual Fellowship 2016-2017. MD is founded by F.R.I.A. (Fonds pour la Recherche dans l’Industrie et l’Agriculture) Fellowship.

## References

1. Vorster, A. P. & Born, J. Sleep and memory in mammals, birds and invertebrates. Neurosci. Biobehav. Rev. 50, 103–119 (2015).

2. Rasch, B. & Born, J. About sleep’s role in memory. Physiol. Rev. 93, 681–766 (2013).

3. Peigneux, P. Neuroimaging studies of sleep and memory in humans. Curr. Top. Behav. Neurosci. 25, 239–268 (2015).

4. Peigneux, P. et al. Are Spatial Memories Strengthened in the Human Hippocampus during Slow Wave Sleep ? Aufbau • Evaluation • Take Home Message • Fragen. Cell Press 44, 535–545 (2004).

5. Orban, P. et al. Sleep after spatial learning promotes covert reorganization of brain activity. Proc. Natl. Acad. Sci. 103, 7124–7129 (2006).

6. Nguyen, N. D., Tucker, M. A., Stickgold, R. & Wamsley, E. J. Overnight Sleep Enhances Hippocampus-Dependent Aspects of Spatial Memory. Sleep 36, 1051–1057 (2013).

7. Burgess, N., Maguire, E. A. & O’Keefe, J. The human hippocampus and spatial and episodic memory. Neuron 35, 625–641 (2002).

8. Boccia, M., Nemmi, F. & Guariglia, C. Neuropsychology of environmental navigation in humans: Review and meta-analysis of fMRI studies in healthy participants. Neuropsychol. Rev. 24, 236–251 (2014).

9. Epstein, R. A., Patai, E. Z., Julian, J. B. & Spiers, H. J. The cognitive map in humans : spatial navigation and beyond. (2017). doi:10.1038/nn.4656

10. Voss, P., Fortin, M., Corbo, V., Pruessner, J. C. & Lepore, F. Assessment of the caudate nucleus and its relation to route learning in both congenital and late blind individuals. BMC Neurosci. 14, 1 (2013).

11. Epstein, R. A., Patai, E. Z., Julian, J. B. & Spiers, H. J. The cognitive map in humans: Spatial navigation and beyond. Nat. Neurosci. 20, 1504–1513 (2017).

12. Hartley, T., Maguire, E. A., Spiers, H. J. & Burgess, N. The well-worn route and the path less traveled: Distinct neural bases of route following and wayfinding in humans. Neuron (2003). doi:10.1016/S0896-6273(03)00095-3

13. Jacobson, T. K., Gruenbaum, B. F. & Markus, E. J. Extensive training and hippocampus or striatum lesions: Effect on place and response strategies. Physiol. Behav. (2012). doi:10.1016/j.physbeh.2011.09.027

14. Packard, M. G. & Knowlton, B. J. Learning and Memory Functions of the Basal Ganglia. Annu. Rev. Neurosci. (2002). doi:10.1146/annurev.neuro.25.112701.142937

15. Moser, M.-B., Rowland, D. C. & Moser, E. I. Place Cells, Grid Cells, and Memory. Cold Spring Harb. Perspect. Biol. 7, a021808 (2015).

16. Wilson, M. A. & McNaughton, B. L. Reactivation of hippocampal ensemble memories during sleep. Science (80-.). (1994). doi:10.1126/science.8036517

17. Nádasdy, Z., Hirase, H., Czurkó, A., Csicsvari, J. & Buzsáki, G. Replay and time compression of recurring spike sequences in the hippocampus. J. Neurosci. (1999). doi:10.1126/science.1182395

18. Grosmark, A. D., Mizuseki, K., Pastalkova, E., Diba, K. & Buzsáki, G. REM Sleep Reorganizes Hippocampal Excitability. Neuron 75, 1001–1007 (2012).

19. Montgomery, S. M., Sirota, A. & Buzsaki, G. Theta and Gamma Coordination of Hippocampal Networks during Waking and Rapid Eye Movement Sleep. J. Neurosci. 28, 6731–6741 (2008).

20. Maquet, P. et al. Experience-dependent changes in changes in cerebral activation during human REM sleep. Nat. Neurosci. (2000). doi:10.1038/77744

21. Peigneux, P. et al. Learned material content and acquisition level modulate cerebral reactivation during posttraining rapid-eye-movements sleep. Neuroimage (2003). doi:10.1016/S1053-8119(03)00278-7

22. Huber, R., Ghilardi, M. F., Massimini, M. & Tononi, G. Local sleep and learning. Nature (2004). doi:10.1038/nature02663

23. Rasch, B., Büchel, C., Gais, S. & Born, J. Odor cues during slow-wave sleep prompt declarative memory consolidation. Science (80-.). (2007). doi:10.1126/science.1138581

24. Peigneux, P. et al. Are spatial memories strengthened in the human hippocampus during slow wave sleep? Neuron 44, 535–545 (2004).

25. Rauchs, G. et al. Sleep modulates the neural substrates of both spatial and contextual memory consolidation. PLoS One 3, (2008).

26. Vinet, L. & Zhedanov, A. A ‘missing’ family of classical orthogonal polynomials. J. Phys. A Math. Theor. 44, 085201 (2011).

27. Urbain, C. et al. Sleep in children triggers rapid reorganization of memory-related brain processes. Neuroimage (2016). doi:10.1016/j.neuroimage.2016.03.055

28. Maquet, P. et al. Memory processing during human sleep as assessed by functional neuroimaging. Rev Neurol (2003). doi:MDOI-RN-11-2003-159-S11-0035-3787-101019-ART4 [pii]

29. Fischer, S. Motor Memory Consolidation in Sleep Shapes More Effective Neuronal Representations. J. Neurosci. (2005). doi:10.1523/JNEUROSCI.1743-05.2005

30. Debas, K. et al. Off-line consolidation of motor sequence learning results in greater integration within a cortico-striatal functional network. Neuroimage 99, 50–58 (2014).

31. Urbain, C., Galer, S., Van Bogaert, P. & Peigneux, P. Pathophysiology of sleep-dependent memory consolidation processes in children. Int. J. Psychophysiol. 89, 273–283 (2013).

32. Peigneux, P. et al. Offline persistence of memory-related cerebral activity during active wakefulness. PLoS Biol. 4, 647–658 (2006).

33. Kelly, C. & Castellanos, F. X. Strengthening connections: Functional connectivity and brain plasticity. Neuropsychology Review (2014). doi:10.1007/s11065-014-9252-y

34. Woolley, D. G. et al. Virtual water maze learning in human increases functional connectivity between posterior hippocampus and dorsal caudate. Hum. Brain Mapp. 36, 1265–1277 (2015).

35. Keller, T. A. & Just, M. A. Structural and functional neuroplasticity in human learning of spatial routes. Neuroimage 125, 256–266 (2016).

36. Buysse, D. J., Reynolds, C. F., Monk, T. H., Berman, S. R. & Kupfer, D. J. The Pittsburgh sleep quality index: A new instrument for psychiatric practice and research. Psychiatry Res. (1989). doi:10.1016/0165-1781(89)90047-4

37. Horne, J. A. & Ostberg, O. A self-assessment questionnaire to determine morningness-eveningness in human circadian rhythms. Int. J. Chronobiol. (1976). doi:10.1177/0748730405285278

38. Oldfield, R. C. The assessment and analysis of handedness: The Edinburgh inventory. Neuropsychologia 9, 97–113 (1971).

39. Ellis, B. W. et al. The St. Mary’s Hospital sleep questionnaire: a study of reliability. Sleep 4, 93–97 (1981).

40. Rauchs, G. et al. Partially segregated neural networks for spatial and contextual memory in virtual navigation. Hippocampus 18, 503–518 (2008).

41. Hutton, C. et al. Image Distortion Correction in fMRI: A Quantitative Evaluation. Neuroimage 16, 217–240 (2002).

42. Friston, K. J., Stephan, K. E., Lund, T. E., Morcom, A. & Kiebel, S. Mixed-effects and fMRI studies. Neuroimage 24, 244–252 (2005).

43. Brown, T. I., Whiteman, A. S., Aselcioglu, I. & Stern, C. E. Structural Differences in Hippocampal and Prefrontal Gray Matter Volume Support Flexible Context-Dependent Navigation Ability. 34, 2314–2320 (2014).

44. Robin, J. et al. Functional connectivity of hippocampal and prefrontal networks during episodic and spatial memory based on real-world environments. Hippocampus 25, 81–93 (2015).

45. Reagh, Z. M. & Yassa, M. A. Repetition strengthens target recognition but impairs similar lure discrimination : evidence for trace competition. 342–346 (2014).

46. Tzourio-Mazoyer, N. et al. Automated Anatomical Labeling of Activations in SPM Using a Macroscopic Anatomical Parcellation of the MNI MRI Single-Subject Brain. Neuroimage 15, 273–289 (2002).

47. Yan Chao-Gan and Zang Yu-Feng. DPARSF: a MATLAB toolbox for “pipeline” data analysis of resting-state fMRI. Front. Syst. Neurosci. 4, 1–7 (2010).

48. Friston, K. J., Williams, S., Howard, R., Frackowiak, R. S. J. & Turner, R. Movement-Related effects in fMRI time-series. Magn. Reson. Med. 35, 346–355 (1996).

49. Whitfield-Gabrieli, S. & Nieto-Castanon, A. *Conn* : A Functional Connectivity Toolbox for Correlated and Anticorrelated Brain Networks. Brain Connect. 2, 125–141 (2012).

50. Behzadi, Y., Restom, K., Liau, J. & Liu, T. T. A component based noise correction method (CompCor) for BOLD and perfusion based fMRI. Neuroimage 37, 90–101 (2007).

51. Deliens, G. & Peigneux, P. One night of sleep is insufficient to achieve sleep-to-forget emotional decontextualisation processes. Cogn. Emot. (2014). doi:10.1080/02699931.2013.844105

52. Jensen, O. et al. Declarative memory consolidation in humans: A prospective functional magnetic resonance imaging study. Proc. Natl. Acad. Sci. (2006). doi:10.1073/pnas.0507774103

53. Haier, R. J., Karama, S., Leyba, L. & Jung, R. E. MRI assessment of cortical thickness and functional activity changes in adolescent girls following three months of practice on a visual-spatial task. 7, (2009).

54. Rajah, M. N., Languay, R. & Grady, C. L. Age-Related Changes in Right Middle Frontal Gyrus Volume Correlate with Altered Episodic Retrieval Activity. J. Neurosci. 31, 17941–17954 (2011).

55. Guterstam, A. et al. Posterior Cingulate Cortex Integrates the Senses of Self-Location and Body Ownership Article Posterior Cingulate Cortex Integrates the Senses of Self-Location and Body Ownership. 1416–1425 (2015). doi:10.1016/j.cub.2015.03.059

56. Nau, M., Navarro Schröder, T., Bellmund, J. L. S. & Doeller, C. F. Hexadirectional coding of visual space in human entorhinal cortex. Nat. Neurosci. 21, 188–190 (2018).

57. Ohnishi, T., Matsuda, H., Hirakata, M. & Ugawa, Y. Navigation ability dependent neural activation in the human brain: An fMRI study. Neurosci. Res. 55, 361–369 (2006).

58. Wong, C. W. et al. Resting-state fMRI activity predicts unsupervised learning and memory in an immersive virtual reality environment. PLoS One 9, (2014).

59. Boccia, M., Sulpizio, V., Nemmi, F., Guariglia, C. & Galati, G. Direct and indirect parieto-medial temporal pathways for spatial navigation in humans: evidence from resting-state functional connectivity. Brain Struct. Funct. 222, 1945–1957 (2017).

60. Hirshhorn, M., Grady, C., Rosenbaum, R. S., Winocur, G. & Moscovitch, M. The Hippocampus Is Involved in Mental Navigation for a Recently Learned, but Not a Highly Familiar Environment : A Longitudinal fMRI Study. 852, 842–852 (2012).

61. Long, X. & Zhang, S.-J. A novel somatosensory spatial navigation system outside the hippocampal formation. doi:10.1101/473090

62. Nguyen, H. M., Matsumoto, J., Tran, A. H., Ono, T. & Nishijo, H. sLORETA current source density analysis of evoked potentials for spatial updating in a virtual navigation task. Front. Behav. Neurosci. 8, 66 (2014).

63. Nelson, A. J. D., Powell, A. L., Holmes, J. D., Vann, S. D. & Aggleton, J. P. What does spatial alternation tell us about retrosplenial cortex function? Front. Behav. Neurosci. 9, 1–15 (2015).

64. Powell, A. L. et al. The rat retrosplenial cortex as a link for frontal functions : A lesion analysis. Behav. Brain Res. 335, 88–102 (2017).

65. Fryer, S. L. et al. Relating intrinsic low-frequency BOLD cortical oscillations to cognition in schizophrenia. Neuropsychopharmacology 40, 2705 (2015).

66. Ren, W., Li, R., Zheng, Z. & Li, J. Neural correlates of associative memory in the elderly: a resting-state functional MRI study. Biomed Res. Int. 2015, (2015).

67. Cavanna, A. E. & Trimble, M. R. The precuneus: A review of its functional anatomy and behavioural correlates. Brain (2006). doi:10.1093/brain/awl004

68. Bubb, E. J., Metzler-Baddeley, C. & Aggleton, J. P. The cingulum bundle: Anatomy, function, and dysfunction. Neurosci. Biobehav. Rev. 92, 104–127 (2018).

69. Epstein, R. & Kanwisher, N. A cortical representation of the local visual environment. Nature 392, 598–601 (1998).

70. Hagewoud, R. et al. A Time for Learning and a Time for Sleep : The Effect of Sleep Deprivation on Contextual Fear Conditioning at Different Times of the Day. 2–9

